# Adaptive collective motions: a hybrid method to improve conformational sampling with molecular dynamics and normal modes

**DOI:** 10.1101/2022.11.29.517349

**Authors:** Pedro T. Resende-Lara, Maurício G. S. Costa, Balint Dudas, David Perahia

## Abstract

Protein function is closely related to its structure and dynamics. Due to its large number of degrees of freedom, proteins adopt a large number of conformations, which describe a highly complex potential energy landscape. Considering the huge ensemble of conformations in dynamic equilibrium in solution, detailed investigation of proteins dynamics is extremely costly. Therefore, a significant number of different methods have emerged in order to improve the conformational sampling of biomolecules. One of these methods is Molecular Dynamics with excited Normal Modes (MDeNM) in which normal modes are used as collective variables in molecular dynamics. Here, we present a new implementation of the MDeNM method that allows a continuously controlled kinetic excitation energy in the normal mode space, while taking into account the natural constraints imposed either by the structure or the environment. These implementations prevent unphysical structural distortions. We tested the new approach on bacteriophage’s T4 lysozyme, Gallus gallus hen egg-white lysozyme and Staphylococcus aureus membrane-bound transglycosylase. Our results showed that the new approach outperformed free MD sampling and preserved the structural features comparatively to the original MDeNM approach. We also observed that by adaptively changing the excitation direction during calculations, proteins follow new transition paths preventing structural distortions.

## 1 INTRODUCTION

The diversity of protein structures and dynamics determine their wide range of functions in organisms as diverse as enzymatic catalysis, signal transduction, protein-ligand, and protein-protein interactions^1,2^, flow control of molecules in and out of the cell, and the control of the cell phenotype^3–6^. The time scales of protein functional internal dynamics generally range from femto to milliseconds covering a wide spectrum of movements from local to collective leading to a huge number of conformations in solution. Their equilibrium distributions on the free energy landscape or the potential energy surface (PES) depend on a particular set of conditions such as temperature, pressure, solvation, presence of ligand etc. Changing these conditions shift the relative populations of states and the transition kinetics between them^7,8^. The pathways derived from the landscapes remain essential to the study of many processes such as protein folding and binding, and ultimately, its function^9,10^; they open the way to establish a relationship between thermodynamics and kinetics, which broadens the structure-function paradigm. Several studies have established such a link using Markov State Models^11–13^, but their applications remain still challenging due to the dimensionality of the energy landscape. They depend on the development of methods to improve or accelerate conformational sampling in biomolecular simulations. A whole class of methods consist of using biased molecular dynamics (MD) simulations such as increased temperatures or modified potential energy functions^14^. Among them the most popular are replica-exchange molecular dynamics^15^, conformational flooding^16^, hyperdynamics^17^, and accelerated molecular dynamics^18^. However, these methods require designing a valid and effective bias potential, with extensive validation. Otherwise, an artificially flat energy surface will be explored.

Another class of methods are capable of performing enhanced sampling without potential bias or loss of detail using collective variables to improve sampling. Methods such as metadynamics^19^, steered molecular dynamics^20^, temperature-accelerated molecular dynamics^21^, amplified collective motions^22^ and umbrella sampling^23^ generally use algorithms that induce the structure to always visit a novel configuration in the conformational space. Nonetheless, any method may fail in properly describing the energy landscape if the collective variables are chosen wrongly or incompletely.

There are also methods that consists of subdividing the total exploration space into subspaces corresponding to different time scales for the movements, thus reducing the extent of the PES exploration. This dimensionality reduction, apart from those used in coarse-grained approaches, explicitly consists of the determination of collective movements taking into account all the atomic degrees of freedom. The most commonly used Principal Components Analysis (PCA) and Normal Mode Analysis (NMA) or Elastic Network Model (ENM), provide a convenient way to define such subspaces, each of these methods having albeit their own limits. One of the major drawbacks of PCA is that the preferential directions of the movements of greater amplitude are closely dependent of the initial conditions of the simulation and its duration, and very long simulation times might be necessary to characterize statistically robust directions and collect all of them. On the other hand, the NMA (or ENM) is a straightforward method to obtain the set of low-frequency directions. One caveat is however that the Normal Mode (NM) vectors are obtained under vacuum conditions with a harmonic approximation of the potential energy function, limiting their validity. But many studies have shown that the lowest frequency directions obtained by NMA although they are valid for small displacements, within the framework of the harmonic approximation, describe quite well the changes observed experimentally at greater distances^24^. Moving at greater distances along these NM directions implies that large anharmonic couplings would occur.

Many methods have been developed using either energy minimizations or MD simulations during the exploration of NM space to take them into account. The simplest method among several other approaches using collective variables is umbrella potentials along the NM directions. A CHARMM implementation called VMOD^25^ was designed for exactly this purpose. Other approaches have been developed, and widely used, such as ClustENM^26^ and CoMD^27^. These methods uses modes predicted by ENM or Anisotropic Network Models (ANM) to explore the conformational space and then interactively deform the structure along these modes by using energy minimization for the former or molecular dynamics in the latter. At the end of each round, the modes are recomputed to be applied in a new iteration round. We have also developed a method that couples local and global motions provided by MD and NMA, respectively, to achieve a better exploration of the conformational space^28–30^.

In the Molecular Dynamics with excited Normal Modes (MDeNM) method, a given NM (or a combination of a NM set) is assigned as additional atomic velocities to the MD simulation and then the system is kinetically excited along this direction to improve the sampling. Usually, all these approaches have been applied to the exploration of the very low-frequency subspace, however higher frequencies which are more localized in nature corresponding to movements of different regions of the macromolecule might also be of interest. An important issue that arises from the use of NM vectors is that they describe rectilinear motions so that domain rotational motions or extensive structural deformations cannot be accounted for easily giving rise to large anharmonic couplings which penalize conformational sampling. At a certain limit, they cease to be representative. NM calculations using dihedral angles are an alternative to overcome such a linearity problem; a comparative study has shown that larger atomic fluctuations are obtained in dihedral space than in Cartesian space^31^, but the NMA in the dihedral space is difficult to apply to whole macromolecules and their complexes considering all atoms. In Cartesian space, a viable alternative is to update the NM vectors during the sampling process^26,32^ to take into account such non-linearity but this may add supplementary computations and other disadvantages such as the discontinuities that occur from sudden changes in NM vectors.

In this article, we present an alternative approach that consists of modifying and adapting the direction of the NM vectors during the displacements such that the tensions resulting from the inherent internal non-linearities and of the environment are reduced allowing greater distances to be explored with minimal stress. We called this adaptive collective motions. We developed this approach within the framework of MDeNM method in which a linear combination of initial modes calculated in the vacuum is used to propagate the movements to larger distances within a complete system comprising the entire environment (water, membrane, etc). We show how small adaptive modifications in collective movement directions allow a more extensive exploration of the conformational space. This leads to further structural changes which would not have been possible if only the initial unchanged NM vectors were considered. Furthermore, we demonstrate that the direction changes take place when low and medium frequency motions are intrinsically activated during the trajectories. We tested this method for different molecular systems obtaining an improved conformational sampling and compared them to those with no directional corrections. The systems considered are the bacteriophage’s T4 lyzozyme (T4L), human calmodulin (CaM) and the *Staphylococcus aureus* monofunctional transglycosylase (MTG).

T4L is a lysozyme that degrades host peptidoglycans, leading to the rupture of the host cell wall and consequently release of the mature viral particles^33^. For many years, it has been used as a model for studying the factors that determine the structure and stability of proteins^34^. It is composed of 164 residues arranged into two domains (N and C) and connected by a long α-helix with the active site cleft (E11 and D20) located at the interface between the two domains^35,36^. The majority of configurations adopted by T4L in experimentally solved structures are related to the closed form (that is similar to the peptidoglycan-bound conformation). However, a more open structure is thought to be required for ligand binding. Indeed, Goto et al. observed structures in which the catalytic cleft is approximately 17° more open than the crystal 3LZM^36^, and Zhang et al. showed that structures with different crystal environments display a range of over 50° in the hinge-bending angle^37^.

Calmodulin is a Ca^2+^-binding protein that modulates the activity of a large number of partners, such as protein kinases, NAD kinases, phosphodiesterases, calcium pumps, and G proteins among others^38^. CaM is composed of two domains formed by the EF-hand helix-loop-helix motifs, which are separated by an interdomain linker. In the calcium-free state (apo-CaM), these domains are collapsed together, assuming a closed conformation where the linker is partially to fully disordered^39,40^. When bound to Ca^2+^ ions (Ca^2+^-CaM), the domains separate from each other and expose a hydrophobic cleft that increases the CaM binding affinity for a plethora of partners^39^. Although the interdomain linker can be seen as an α-helix, this region presents high flexibility and it this feature is of great importance to the wide interdomain orientations responsible to the binding to the most diverse proteins^40,41^.

MTG is a membrane-bound peptidoglycan polymerase that catalyzes glycan chain elongation, essential to the synthesis of bacterial cell wall^42^. The transmembrane (TM) helix plays a crucial role in activity, by contributing to the hydrophobic interaction and maintaining the proper orientation at the membrane for enzymatic catalysis^43^ leading in the approach of the jaw and head subdomains of MTG closer to each other, allowing the formation of the lipid binding pocket^42^. Although there are not many structures available, the solved ones (apo and different ligands bound) reveal a similar orientation within the lipid layer, presenting a good agreement with an *E. coli* analog, the penicillin-binding protein 1b^43^. The essential role of MTG to bacterial survival makes of it a target to the development of antibiotics. Nonetheless, since the mechanism of transglycosylation reaction has been recently unveiled, the design of new structure-based antibiotics targeting this enzyme is still an ongoing effort.

In the following section we present a summary of the MDeNM method, followed by the presentation of the new MDeNM implementations (energy injection control and updating of excitation directions). Section 3 give applications and analysis of the results for the illustrative examples mentioned above.

## 2 METHODS

Molecular Dynamics with excited Normal Modes^28^ was developed to efficiently explore the dynamic equilibrium of macromolecules without introducing any bias in the potential terms. This method consists of periodically applying an additional energy along the direction of a previously selected low frequency normal mode (or a linear combination of a normal modes set) during the molecular dynamics calculation. This additional energy determines the velocity that is added to the current system promoting a motion along a random linear combination of modes. Through this combination, it is possible to obtain a larger exploration of the system’s conformational space, when compared to the high computational cost of regular molecular dynamics. This approach allows the simulation of large time scale movements, such as allosteric transitions of domains. The advantage of this method is the absence of constraints or potential biasing. However, structural and environmental effects during the conformational space exploration may be neglected if the kinetic energy input is too high. In spite of the advances brought by MDeNM, the cumulative character of the energy injection may cause structural distortions along the simulation if the parameters are not efficiently tested. In addition, by exploring the direction defined by the normal modes combination, effects caused by the structure itself or the medium on the directionality of the motion may be underestimated or neglected. Herein, we present novel developments in MDeNM technique to avoid these setbacks. The new MDeNM implementations are depicted at the flowchart on figure 1.

**Figure 1.**
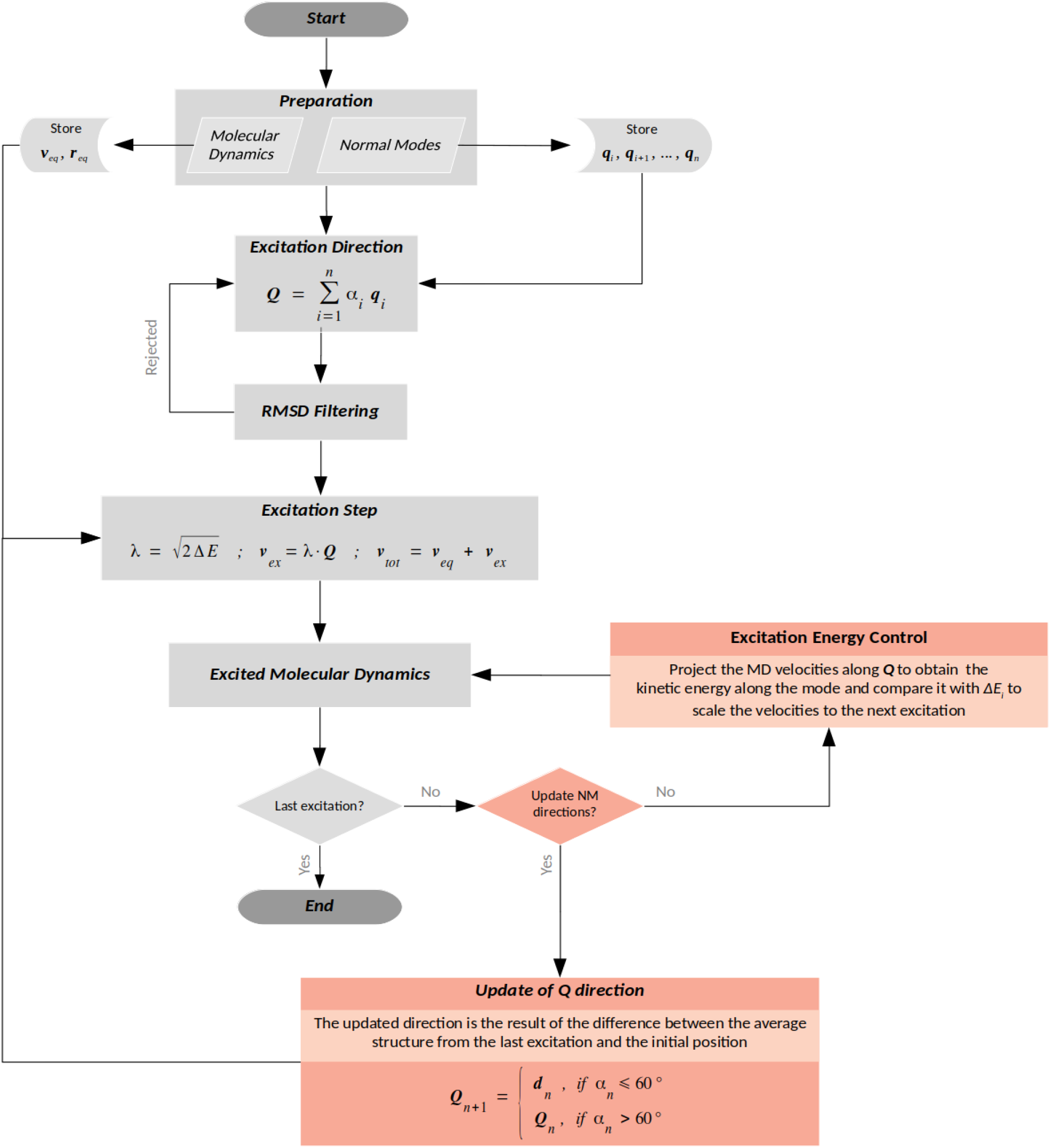
MDeNM flowchart with the new implementations. The preparation step consists of running a short MD and computing the normal modes of the minimized structure resulting in the initial positions (***r***_*eq*_) and velocities (***v***_*eq*_) as well as the directions to further excite the system (***q***_*1*…*n*_), respectively. Afterwards, the excitation vector (***Q***) is created from a random linear combination of different NMs. In a multi-replica procedure, each replica has a different ***Q*** vector. The vectors are then compared to each other in order to retain only those sufficiently different, avoiding unnecessary computation (RMSD filtering step). Once the vector is retained, it will be multiplied by a scalar (*λ*) that yields the additional energy (*ΔE*) set by the user resulting in the excitation velocities (***v***_*ex*_), that are ultimately added to the previous ones (***v***_*eq*_). The simulation is then started. After each excitation, the energy along the vector ***Q*** is evaluated and, if necessary, re-scaled to the amount chosen by the user. Eventually, the real displacement direction of the system is compared to the theoretical and updated if there is a 60° deviation between them.

### 2.1 Monitoring Kinetic Energy Along the Excitation Vector

In order to keep the structure in a permanently excited state during the simulations, we developed a new algorithm that constantly evaluates and corrects the kinetic energy injection along the excitation direction. This procedure is required since the additional kinetic energy has a fast dissipation^28^. Therefore, at the end of each excitation the final velocity along the excited direction is evaluated and re-scaled if necessary so as to maintain a constant kinetic energy level. Thus, the system is kept in a continuous excited state, allowing effective small energy supply. The energy injection control is done by projecting the velocities computed during the excitation onto vector ***Q*** and obtaining the kinetic energy corresponding to it, as follows:

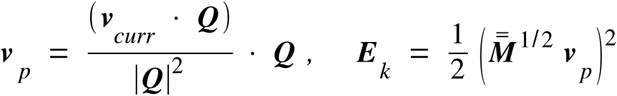

where ***v***_*curr*_ is the current velocity vector, |***Q***|^*2*^ is the norm of vector ***Q*** and 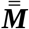 is the mass matrix. As ***v***_*exc*_ is a constant excitation energy input defined by the user, and knowing the current ***v***_*p*_ by the above equation, one can obtain the new excitation velocities to inject as defined by ***v***_*new*_ *=* ***v***_*curr*_ *+ (****v***_*exc*_ *–* ***v***_*p*_). Detailed comparison between the energy input along the simulation in original and ceMDeNM is illustrated in the supp. fig. 1.

### 2.2 Update of Excitation Direction

Considering that vector ***Q***_*n*_ is obtained from the initial structure, its direction is intrinsically dependent of this initial position. This means that the identity of the motion noticeably decreases after successive excitations due to anharmonic effects. Thus, it is necessary to change these directions all along the simulation and, consequently, prevent structural distortions produced by the excitation along the deteriorated ***Q***_*n*_ vector. This procedure allows the system to adaptively find a relaxed path to follow during the next MDeNM excitation steps. To this end, we developed an algorithm that writes an adapted excitation direction (***Q***_*n+1*_) based on the trajectory evolution during the MDeNM run. The correction is done based on two parameters: one related to the effective displacement of the structure along the excitation vector ***Q***_*n*_, (*ℓ*); and the other considering the relative deviation of the effective displacement vector with respect to vector ***Q***_*n*_, (*cos α*). The unit vector ***d***_*n*_ is computed accordingly to the following equation: 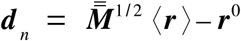, where ***d***_*n*_ is the mass-weighted difference vector between the average position (*‹****r****›*) of the last excitation along vector ***Q***_*n*_ and the initial position of this segment (***r***^*0*^). In addition, the scalar *cos α* represents the projection of ***d***_*n*_ onto ***Q***_*n*_, ***d***_*n*_*/* | ***d***_*n*_| · ***Q***_*n*_ = | ***Q***_*n*_ |cos α=cos μ.

The excitation vector is changed as soon as the trajectory reaches a *ℓ* threshold distance along ***Q***_*n*_ and *cos α* is lower than a reference value (0.5, as shown in Supp. Materials). Otherwise, the simulation is resumed with the previous accepted excitation directions (***Q***_*n*_). The relationship between *ℓ* and *cos α* is discussed in the supplementary text as well as supp. fig. 2. Then, the

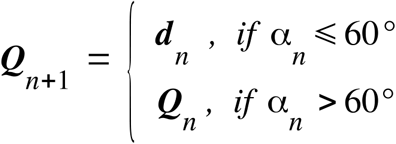

corrected vector is defined as:

Figure 2 illustrates the excitation vector correction along MDeNM simulations.

**Figure 2.**
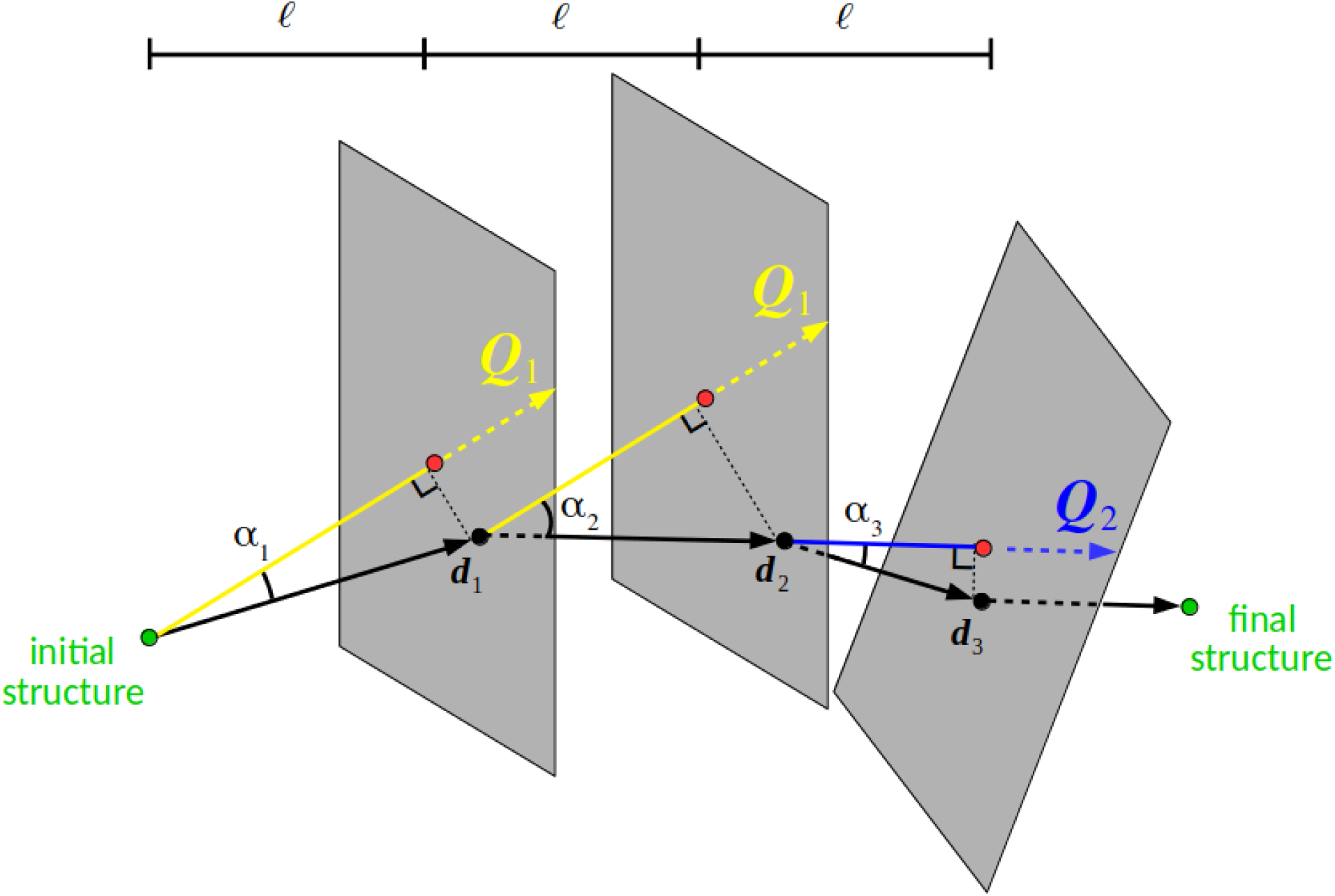
Adaptive correction of excitation vector during MDeNM simulations. The vectors *Q*_*1*_ and *Q*_*2*_ correspond to excitation directions and *d*_*1-3*_ to the effective mean displacement vectors during the corresponding excitation. After reaching a given MRMS displacement threshold (*ℓ*) along the excitation direction, the program evaluates the need to change the excitation direction (evaluations are shown as gray planes). Excitation directions are updated during the simulation if *cos α*_*n*_ (being *α*_*n*_ the angle formed between the vectors *d*_*n*_ and *Q*_*n*_) is equal or lower than 0.5, that is, if the real displacement vector presents a deviation larger than 60° from the theoretic one. The variables *ℓ* and *cos α*_*n*_ represent, respectively, the relative distance and orientation in respect with the original position and excitation vector. In this image, the first evaluation results on *cos α*_*1*_ ≥*0*.*5*. Therefore, the excitation direction remained the same as the previous excitation (*Q*_*1*_). Afterwards, *cos α*_*2*_ *< 0*.*5* in the following evaluation. Thus, a new excitation vector is written (*Q*_*2*_ *= d*_*2*_). By doing this, one can enable the exploration of new directions that take into account the effects of the medium on the protein motions.

The direction change must take place after a displacement large enough to be greater than the normal fluctuations, otherwise the simulation precision is lost. Likewise, it is necessary that the change of orientation also be sufficiently large to proceed with the update. If the change of orientation is large but the displacement is small, and vice versa, the update is not done because there is too much uncertainty due to fluctuations.

### 2.3 Probed systems and simulation details

#### 2.3.1 Atomic Coordinates and Experimental Datasets

The atomic coordinates of bacteriophage’s T4 lysozyme (T4L, PDB id: 178L^37^), human calmodulin (CaM, PDB id: 1CLL^44^) and Staphylococcus aureus membrane-bound transglycosylase (MTG, PDB id: 3VMQ^43^) were obtained from the Protein Data Bank and employed in the validation set of simulations presented in this study. The few residues gap in the structure 3VMQ was modeled with MODELLER 9.19^45^. Co-crystalized water molecules were maintained in final models. Hydrogen atoms were added/discarded to all structures in order to change the protonation state of residues, by using PROPKA3^46^ corresponding to a pH of 7.4.

#### 2.3.2 Molecular Dynamics

T4L and CaM molecular dynamics systems were built by CHARMM-GUI input generator^47,48^. Molecular dynamics were performed using CHARMM c41b1 and the CHARMM^49,50^ force field parameter set 36^51^ with explicit TIP3 water molecules, using periodic boundary conditions. The PME algorithm was applied to treat electrostatic interactions^52^. The protein was kept at least 10Å apart from the boundaries of an octahedral box. Chloride ions were added for charge balancing. The NVT equilibration dynamics was carried with the Velocity Verlet integrator with an integration time of 1fs during 25ps. Temperature was kept constant by using the Nosé-Hoover algorithm^53,54^. The SHAKE^55^ algorithm was used to fix water and protein bonds. Protein heavy atoms were harmonically restrained with a force constant of 2.0 and 0.2 *m*_*i*_ kcal/mol^-1^Å^-2^ (where *m*_*i*_ is the atomic mass of the atom *i*), for backbone and sidechain heavy atoms, respectively, in order to avoid artificial structural distortions. Then, an unconstrained NPT production MD was carried with the Leapfrog Verlet integrator for 125ns, with temperature and pressure kept constant by using the Nosé-Hoover algorithm. The reference temperature was set to 303.15K and the pressure to 1 atm.

We used the CHARMM-GUI membrane builder^56–58^ to generate MTG molecular dynamics inputs. Firstly, we used GROMACS 5.0.5^59^ and CHARMM force field parameter set 36 in order to equilibrate MTG in the membrane. The insertion of the protein into the membrane was done following OPM orientation^60^. The bilayer composition was 43% palmitoyl-oleoyl-phosphatidylglycerol (POPG), 30% palmitoyl-linoleoyl-phosphatidylglycerol (PLPG), 22% 1,1′,2,2′-tetraoleoyl cardiolipin (TOCL1) and 5% of phosphatidic acid (POPA), according to the data available in ref.^61^. Explicit TIP3 water molecules and potassium ions were added to the tetragonal box. Protein was kept at least 17.5Å apart from the box boundaries. After 5000 steps of steepest descent (SD) energy minimization, we based the equilibration protocol as described in ref.^57^ (Table 1) and a NPT production dynamics of 125ns was carried out removing all constraints using the Leapfrog Verlet integrator, with temperature and pressure kept constant and fixed to 303.15K and 1 atm, respectively, by using the Nosé-Hoover algorithm.

**Table 1.** Detailed information on each equilibration step.

| Step | Ensemble<sup>1</sup> | Timesteps | Equilibration Time | Force constants for Harmonic Restraints<sup>2</sup> |  |  |  |  |
| --- | --- | --- | --- | --- | --- | --- | --- | --- |
|  |  |  |  | Protein Backbone<sup>3</sup> | Protein Sidechain<sup>3</sup> | Water<sup>4</sup> | Lipids<sup>5</sup> | Ions<sup>3</sup> |
| 1 | NVT | 1 fs | 25ps | 10.0 | 5.0 | 2.5 | 2.5 | 10.0 |
| 2 | NVT | 1 fs | 25ps | 5.0 | 2.5 | 2.5 | 2.5 | 0.0 |
| 3 | NPT | 2 fs | 25ps | 2.5 | 1.0 | 1.0 | 1.0 | 0.0 |
| 4 | NPT | 2 fs | 100ps | 1.0 | 0.5 | 0.5 | 0.5 | 0.0 |
| 5 | NPT | 2 fs | 100ps | 0.5 | 0.1 | 0.1 | 0.1 | 0.0 |
| 6 | NPT | 2 fs | 100ps | 0.1 | 0.0 | 0.0 | 0.0 | 0.0 |

<sup>1</sup> NVT stands for constant volume and temperature, and NPT for constant pressure and temperature. <sup>2</sup> Force constants are in kcal/(mol×Å<sup>2</sup>). <sup>3</sup> Positional harmonic restraints. <sup>4</sup> Harmonic restraints to keep water molecules away from the membrane hydrophobic region. <sup>5</sup> Harmonic restraints to keep the lipid tail in $-5\text{Å} > Z > 5\text{Å}$ , and lipid head groups close to the membrane surface.

#### 2.3.3 Normal Modes Calculations

Energy minimization and NM calculations were carried out using CHARMM c41b1^50^ and CHARMM force field parameter set 36^51^. Van der Waals interactions were calculated up to 10Å, being approximated until 12Å by using a switching function. Electrostatic interactions were calculated up to 10Å. We used the steepest descent (SD) and the Conjugated Gradients (CG) methods to minimize the energy of the structures. This was followed by another energy minimization using the Adopted-Basis Newton Raphson (ABNR) algorithm. Harmonic restraints were applied during 5×10^4^ SD steps, being progressively decreased from 250 to 0 kcal/mol^-1^Å^-2^. Afterwards, we minimized the system with 10^6^ steps of CG and then applied the ABNR algorithm with no constraints using a convergence criterion of 10^−5^ kcal/mol^-1^Å^-2^ RMS energy gradient. The 200 lowest frequency normal modes for all atoms (dismissing the 6 modes related to rotation and translation around the axes) were computed in vacuum using the DIMB^62^ module implemented in CHARMM. A distance dependent dielectric constant *(ε=2r*_*ij*_) was employed to treat electrostatic interactions. NM and atomic fluctuations (root mean square fluctuations – RMSF) were computed with the Vibran CHARMM module.

#### 2.3.4 MDeNM setup

In this study, we have compared the conformational sampling of three different MDeNM applications (original, constant energy input and adaptive MDeNM), free molecular dynamics simulations and the available experimental structures. In the original MDeNM approach, an incremental energy of 5 kcal/mol was added to the system every 2ps. In contrast, the constant energy MDeNM approach (ceMDeNM) injects 0.125 kcal/mol to the system every 0.2ps and maintain this same excitation level during the whole simulation. This differs from the original approach since the relaxation time here is noticeably reduced, so the protein was kept always in an excited state. The excitation in the adaptive MDeNM (aMDeNM) approach was the same as previously described. The difference lies on whether the algorithm update the excitation direction or not based on the conditions explained at the section 2.2.

#### 2.3.5 Relevant internal coordinates

##### 2.3.5.1 T4 lysozyme

We evaluated the T4L hinge bending and torsion motions since they are the most important motions to its dynamics^35,63^. The T4L hinge angle θ can be obtained from the computation of the center of mass of Cα atoms from three regions: (1) residues 19-23 in the N-domain; (2) 68-72 in the middle of the interdomain α-helix and (3) 137-141 in the C-domain. Whereas the torsion angle γ between the domains is composed by the dihedral formed between the center of mass of Cα atoms of: (1) residues 19-23 in the N-domain; (2) residue 61 at the beginning of the interdomain α-helix; (3) residue 79 at the end of the interdomain α-helix and (4) 137-141 in the C-domain.

##### 2.3.5.2 Calmodulin

As described by Aykut et al.^*64*^, the domain torsion and the linker end-to-end distance are two efficient measures to identify different conformational states of CaM. The torsion angle γ between the domains is composed by the dihedral formed between the center of mass of Cα atoms of: (1) residues 8-68 in the N-domain; (2) residue 69 at the beginning of the interdomain linker; (3) residue 91 at the end of the interdomain linker and (4) 92-142 in the C-domain. The end-to-end linker distance is computed as the distance between the Cα atoms of residues 69 and 91.

##### 2.3.5.3 Monofunctional transglycosylase

Considering the importance of the TM helix to the enzymatic activity of MTG, we evaluated its orientation inside the membrane highlighting the bending and the rotation of the helix. The rotation is measured by monitoring the variation, in degrees, of the longitudinal axis along the trajectory while the bending refers to the angle formed Cα atoms of residues R41, I53 and R67. In addition, we observed the sampling of the globular domains independently. We also assessed the Cα-RMSD and the hinge angle θ obtained from the center of mass of Cα atoms as following: (1) residues 67-130 and 168-269 in the head domain; (2) 106 and 167 interconnecting the two globular domains and (3) 107-126 in the jaw domain.

#### 2.3.6 Analytical tools

Visual representations of the proteins and plots were made with PyMOL 2.3.2^65^ and R 3.4.4, respectively.

## 3 RESULTS AND DISCUSSION

In the following, we will compare the structural sampling of free molecular dynamics and three MDeNM applications: original (as published by Costa et al.^28^), constant energy injection and adaptive MDeNM. The systems were chosen to evaluate different features of the MDeNM method: *i)* T4L has its overall dynamics described by two low frequency normal modes, so its dynamics can be easily described; *ii)* Calmodulin has many degrees of freedom and, therefore, it may be intricate to describe its motions; and *iii)* MTG is a transmembrane protein, then, the heterogeneous environment may play a role differently in its motions. We also rationalize the differences between MDeNM applications in the light of vibrational and free energy analysis.

### 3.1 Broader sampling of T4L with MDeNM approaches

To explore the hinge bending and domain twisting, we applied MDeNM along 48 different combinations of normal modes 7 and 8 (see Supp. Material for details), that are responsible for roughly 90% of T4L’s overall dynamics^63^. Starting from an open conformation, we simulated T4L closing motion. After 125ns of free MD we observed a rather poor exploration, where the protein was trapped within a minimum energy well between the states most of the time (fig. 3a). In spite of the relatively large simulation time, not all experimental structures were covered by free MD. By applying MDeNM we observed a great difference in this pattern. The original approach has improved the sampling significantly (fig. 3b). Moreover, both open and closed states were well visited during the 12ns of simulation (48 replicas of 250ps each) as shown by the hot colors on the density map. This is consistent with the previous works where it was shown the ability of easily shift between different protein macrostates using MDeNM^28–30^. It is important to point out that the conformations most visited were the ones with the lowest potential energy in comparison to the minimized structure (fig. 3c). When using constant energy injections, we observed an even broader sampling than the previous ones (fig. 3d). The motions were exaggerated and presented some level of structural distortions such as the slightly loss of secondary structure (supp. fig. 3). Moreover, the closed conformation was less populated and the potential energy of the states visited in this approach was higher the original MDeNM (fig. 3e). This indicates that the system retains more energy to itself in ceMDeNM, implicating in less relaxation and, therefore, crossing larger energy barriers (not necessarily into biologically relevant conformational states). On the other hand, adaptive MDeNM presented very good sampling, densely populating the open and closed macrostates (fig. 3f). To change the excitation directions on-the-fly was shown efficient in avoiding structural deformations as the secondary structure was essentially kept (supp. fig. 3). Similarly to the original approach, the potential energy was lower at the surroundings of the transition path between open to closed conformations (fig. 3g) although we observed wide open and twisted structures with low relative energy as well (right upper corner, fig. 3g).

**Figure 3.**
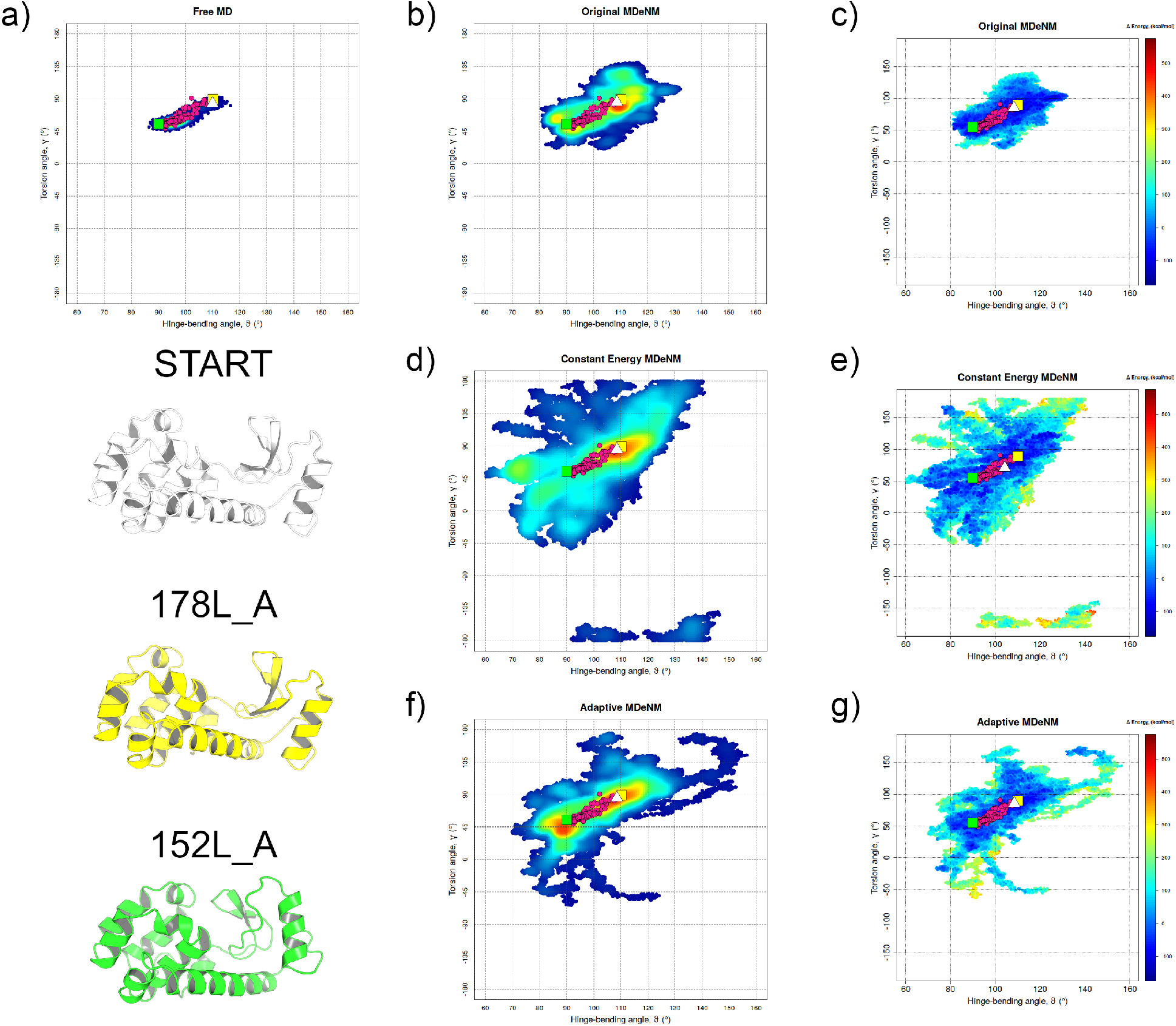
T4L sampling improvement using MDeNM applications. Density scatter-plot demonstrating the sampling of T4L through **a)** free MD; **b)** MDeNM; **d)** ceMDeNM; and **f)** aMDeNM. The least visited regions are shown in blue while the most ones are displayed in red. Potential energy (*E*_*initial*_ *-E*_*current*_) is shown for **c)** MDeNM; **e)** ceMDeNM; and **g)** aMDeNM. Magenta circles represent the experimental structures. White triangle indicates the simulations initial structure. Yellow and green squares indicate an open and closed structures, respectively.

The ceMDeNM and aMDeNM approaches considerably enhanced the sampling compared to the free MD. With less than 20% of computing time, we covered more than 15 times the conformational landscape formed by the chosen internal motions. Our results are compatible to those obtained by Zhang and colleagues using MD accelerated by ANM modes (ACM)^22^ or principal component analysis (ACM-PCA) in order to enhance the sampling^66^. In their approach, ANM or PCA modes are frequently recomputed in order to improve the sampling during the MD simulation without violating the structural constraints of the system. They observed larger motions with their method than to free MD, respecting the overall secondary structure content of T4L. In a recent study^67^, ENM modes are used as geometry constraints to explore large protein motions, enabling the protein to accurately “jump” between macrostates described by the combination of a few modes. The results for T4L showed that the combination of the 3 lowest-frequency modes is required to describe the transition between the PDBs 177L (closed) to 178L (open) states. Here, we presented that the transition between modes 7 and 8 were sufficient to efficiently describe the open-to-close transition.

Considering the update of the excitation direction, the final vector might be quite different from the first one. We observed the activation of different internal motions during the excited trajectory. Figure 4 illustrate this process. As presented at the beginning of this section, we used 48 combinations of modes 7 and 8 to obtain the excitation vectors. Here, we show the projection of the trajectory onto these modes. In the first three panels, only normal mode 7 (hinge-bending motion) was taken into account in order to excite the system. The original MDeNM approach led T4L from the open state to the closed one, where it finds a barrier and stay oscillating around the same conformation (fig. 4a). In contrast, ceMDeNM crossed that barrier and sampled further on the closing motion (fig. 4b), however, showing some level of structural distortion at the endpoint. Both MDeNM and ceMDeNM have not presented any contribution from mode 8 (torsion) during the trajectory (values about 0.6 are due to thermal oscillations). It is interesting to observe that the mode 8 was fully activated during the adaptive approach (fig. 4c). As the trajectory evolves, the mode 7 continuously loses importance to mode 8. Notably, the final displacement along mode 7 was the same that in the original approach. The pathway, however, was quite different. In addition, aMDeNM reached the closed conformation slightly better than the original approach (Cα-RMSD of 1.15Å against 1.21Å). In spite of all 48 aMDeNM replicas account only modes 7 and 8 (fig. 4d), we also observed relevant contributions of modes 9 and 10 (fig. 4e) as well as 11 and 12 (fig. 4f). This indicates that the protein activates multiple internal motions in order to relax the path along the trajectories.

**Figure 4.**
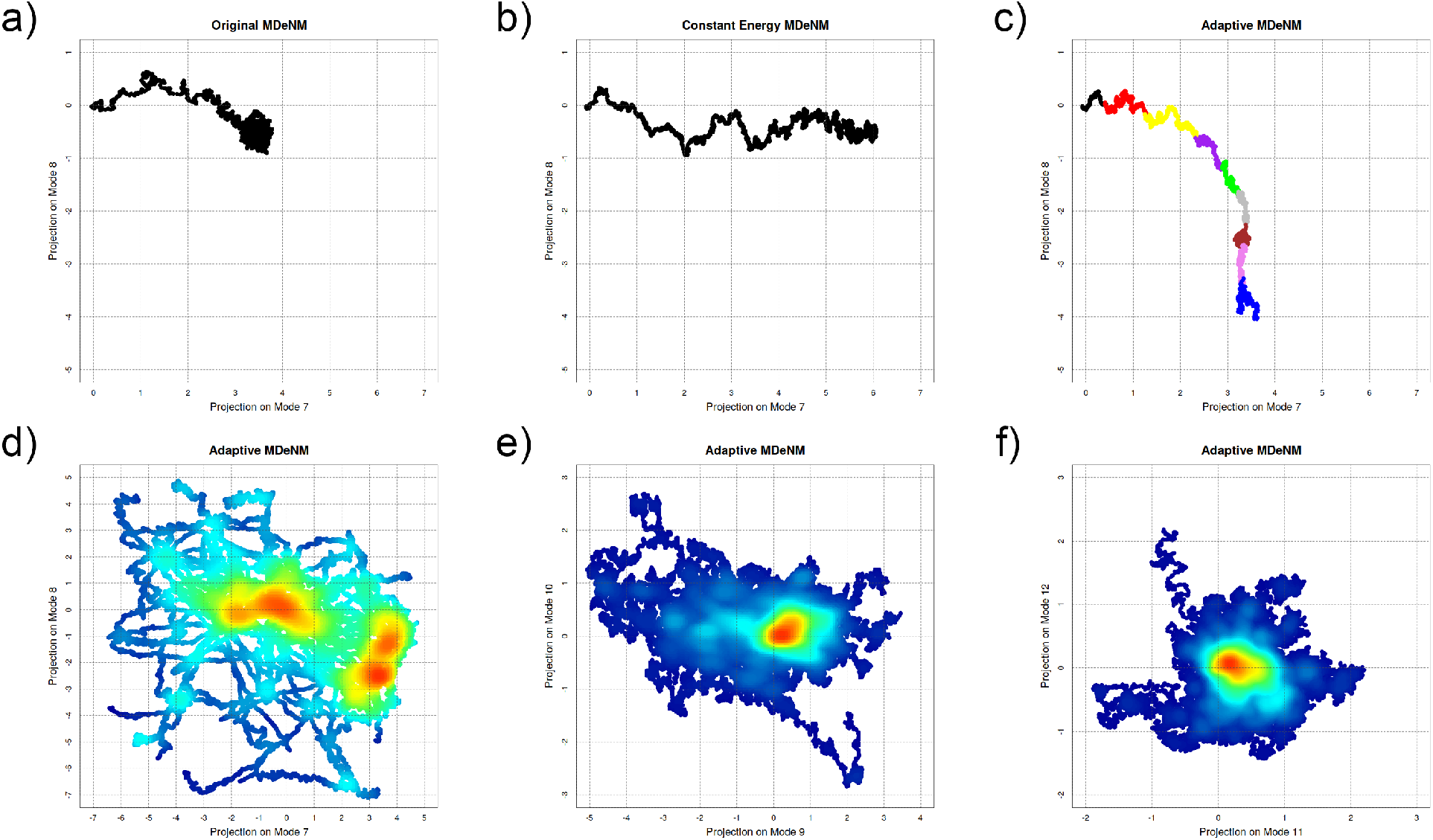
Activation of different internal motions due to the adaptive approach. Using only NM7 to perform the excitation, remarkable differences were observed when projecting the MDeNM trajectories onto normal modes 7 and 8. **a)** In the original MDeNM approach, T4L undergoes a displacement of 3.5Å along the NM7 axis and then it is blocked. **b)** When applying ceMDeNM, the trajectory along NM7 is greatly enhanced, due to the cumulative energy stored by the protein during the simulation. **c)** aMDeNM presented a clear activation of NM8 in its trajectory. After multiple updates of the excitation direction (each color represent a different vector computed during the simulation), the NM7 is almost completely replaced by the NM8. Taking into account the 48 different combinations between NMs 7 and 8, we observe **d)** a very broad sampling of these two NMs, as expected; but also the activation of modes **e)** 9 and 10 as well as **f)** 11 and 12, in a lower degree.

### 3.2 CaM complex motions better represented by new the MDeNM approaches

The binding of Ca^+2^ ions induces significant conformational changes on CaM. This conformational shift depends, among other factors, on the modulation of electrostatic interactions in the highly flexible interdomain linker between the two CaM domains^64^. This modulation lead to the linker assuming a compact or extended conformation, that ultimately favor the binding of several different partners. Here we explore the sampling of CaM by using 48 combinations of NM 7 and 8 and monitoring the reduced degrees of freedom described at section 2.3.5.2.

The initial structure presented an extended conformation with the linker fully structured as α-helix. During 125ns of free MD, CaM mainly fluctuated around this configuration and did not reflected the experimental landscape as shown in fig. 5a. Despite increasing considerably the sampling, MDeNM was not able to reproduce the experimental data either (fig. 5b). The ceMDeNM approach presented a similar pattern, however, it was able to enrich the sampling at a small region near the structure 5WC5_R (right upper corner, fig. 5d), which present the Ca^+2^-CaM complexed with a small-conductance Ca^+2^-activated K^+^ channel. In this conformation, the linker is partially structured and the domains are twisted about 120°. In contrast, aMDeNM displayed an extensive sampling that covered almost all experimental data (fig. 5f). This broader sampling is considerably interesting to observe in a protein with such flexibility, indicating that our method describes very well complex, non-linear motions. Curiously, the region near the structure 5WC5_R was not visited (right upper corner, fig. 5f). Since this region was only visited by the ceMDeNM approach, it could indicate that there is an energy barrier high enough to obstruct the sampling of these conformations. Considering that ceMDeNM stores more energy than the other approaches (due to a smaller relaxation time), the system could overcome that barrier and visit the region. Similarly to that observed for T4L, the regions most densely populated were the ones with lowest potential energy (fig. 5c, 5e and 5g).

**Figure 5.**
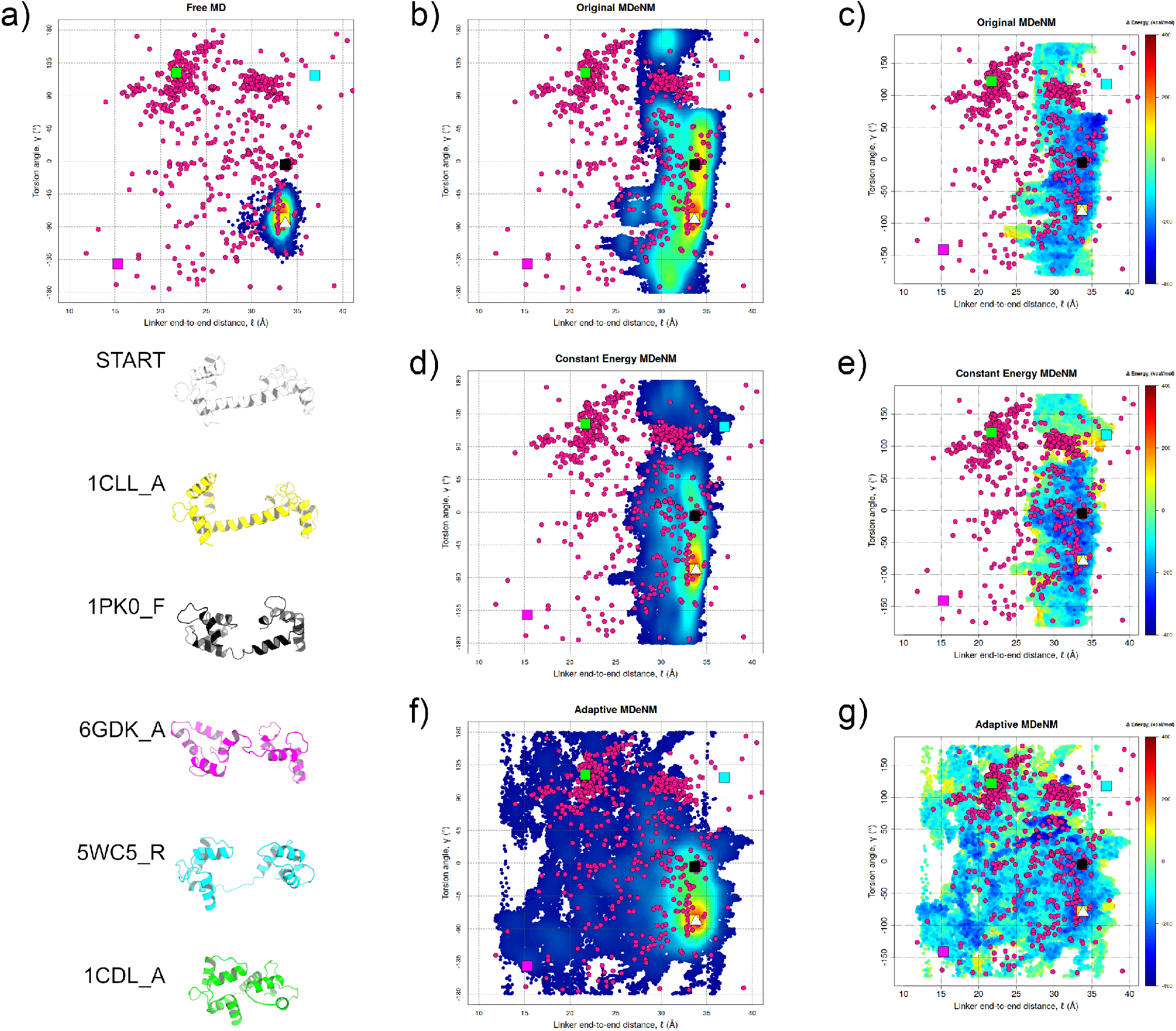
Adaptive MDeNM radically improved CaM sampling. Density scatter-plot demonstrating the sampling of CaM through **a)** free MD; **b)** MDeNM; **d)** ceMDeNM; and **f)** aMDeNM. The least visited regions are shown in blue while the most ones are displayed in red. Potential energy (*E*_*initial*_ *-E*_*current*_) is shown for **c)** MDeNM; **e)** ceMDeNM; and **g)** aMDeNM. Magenta circles represent the experimental structures. White triangle indicates the simulations initial structure. Colored squares indicate experimental structures with diverse conformations, as showed by the cartoon representation.

As previously presented, the highly flexible CaM linker allows the protein to assume a huge plethora of different conformations. This feature is somewhat captured by the broad spanning in the two degrees of freedom presented in fig. 5. However, in the light of such complex motions, we also evaluated the RMSD of our generated structures and compared to some representatives experimental conformers in order to robustly evaluate the conformational sampling of MDeNM approaches. We observed that the our new approaches improved the overlapping between the experimental data and the theoretically generated structures, in particular the aMDeNM (table Table 2). Comparing the probability density function for the different techniques, it is clear that both ceMDeNM and aMDeNM overcome the sampling provided by the original MDeNM and free MD in every scenario (fig. 6). In addition, secondary structure data shows that aMDeNM generated structures are in closer agreement to the experimental data than the other approaches, in spite of some regions keep more residues in α-helix than the observed experimentally (supp. fig. 4).

**Table 2.** Minimal RMSD to selected experimental structures

| Structure | Free MD | MDeNM |  |  |
| --- | --- | --- | --- | --- |
|  |  | Original | Constant Energy | Adaptive |
| **1CLL_A** | 1.748 | 1.739 | 1.547 | **1.321** |
| **1PK0_F** | 5.346 | 7.610 | 3.806 | **3.689** |
| **6GDK_A** | 10.347 | 9.193 | **3.709** | 5.240 |
| **5WC5_R** | 9.634 | 8.962 | 7.526 | **3.586** |
| **1CDL_A** | 12.052 | 10.994 | 8.844 | **4.435** |

**Figure 6.**
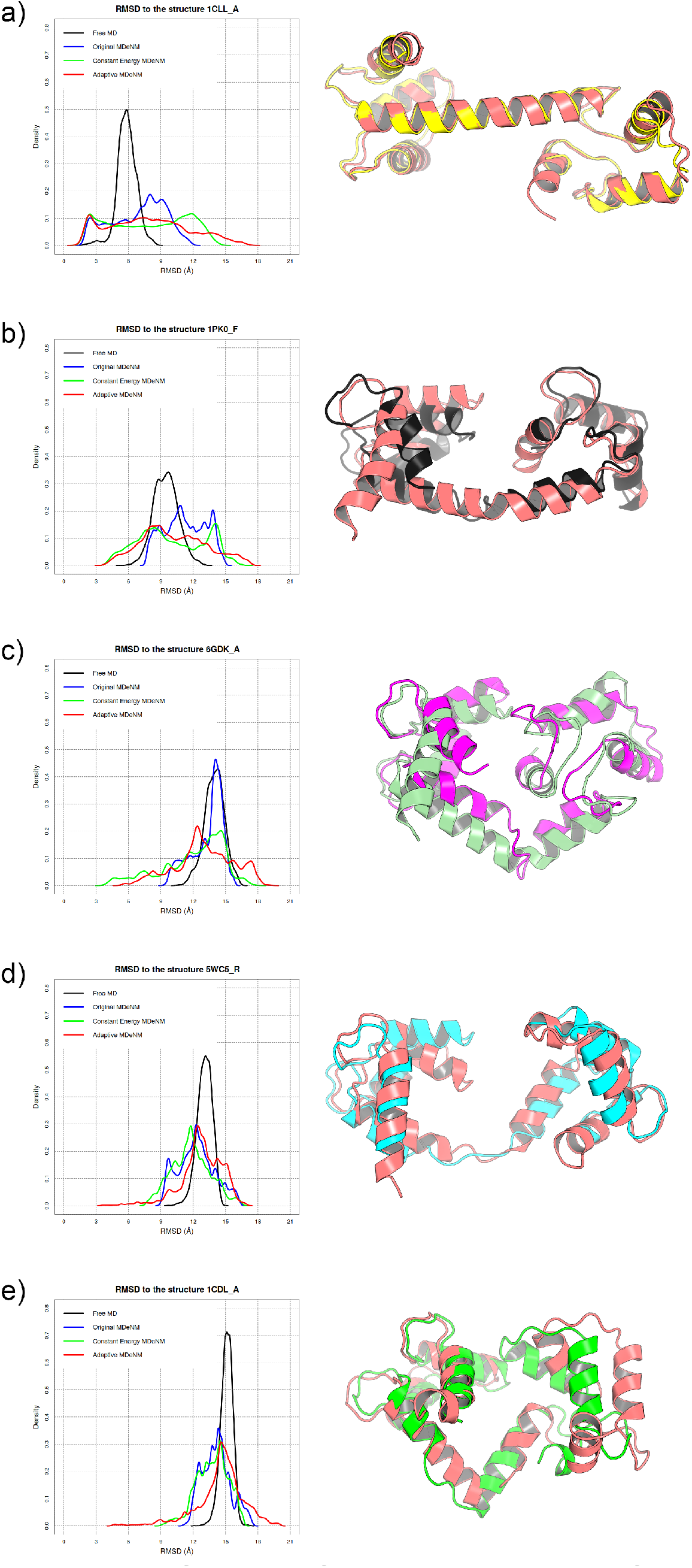
Probability density function of RMSD distribution of selected experimental structures. Both ceMDeNM and aMDeNM displayed broader sampling than free MD and original MDenM when compared to the following experimental structures: **a)** 1CLL_A (yellow) **b)** 1PK0_F (black) **c)** 6GDK_A (magenta) **d)** 5WC5_R (cyan) and **e)** 1CDL_A (green). The pale red cartoons are the minimal RMSD structure generated by aMDeNM whereas the pale green was obtained by ceMDeNM.

The complex motions of CaM were targeted by several enhanced sampling techniques during the past few years. One of these methods combines NMA with an integrated-tempering-sampling molecular simulation (NMA–ITS)^68^. This technique consists in a biased potential MD that uses NMA to drive the motions between two previoulsy known macrostates. We demonstrate comparable results with our technique considering the internal angles of CaM and the overlapping with the experimental data using a method that requires no previous knowledge of the path between the structures. An iterative method to improve protein sampling, ClustENM uses ANM modes to deform the structure and generate the conformational subspace during successive steps and then minimize the generated conformers with an implicit water model^26^. Without the need to recompute the normal modes in every excitation, aMDeNM presented better results, in special for the collapsed conformations. Saldaño et al. present a similar method that uses NMA to generate a realistic ensemble of conformation that describes the protein equilibrium dynamics at a given temperature^32^. This method works both biased by a target structure or unbiased. Our results are consistent with their findings even using only two modes as input instead of ten. The observation that a few middle-frequency modes are required to efficiently reproduce the CaM conformational landscape led to complementary simulations that improved the observed sampling. Nevertheless, we observed that these frequencies were activated during aMDeNM simulations and that the updating of the excitation direction take them into account.

### 3.3 MGT: broader sampling of globular domain and TM helix stable

It has been observed that the TM helix plays a role in the enzymatic activity of MGT, maintaining a proper membrane orientation^43^. The TM helix bending and rotation were assessed to evaluate the protein insertion in the membrane. The bending angle remained between 140° and 180° in all simulations (fig. 7a-c). However, we observed that the in free MD the TM helix presented a rotation of approximately 20° (fig. 7a), much more narrow than ceMDeNM, that reaches rotations up to 60° (fig. 7b). This is consistent to the NM vectors, which present high flexibility in the TM helix. Thus, the excitation artificially exacerbates these motions since the NM are computed in vacuum and, therefore, this region has no restrictions to move. This is clearer when compared to the adaptive approach. Adaptively changing the excitation directions showed the TM much more stable (fig. 7c), considering that the membrane constraints the helix motions in spite of the energy injection. On the other hand, the sampling of MGT globular domains was greatly increased in aMDeNM (fig. 7f) compared to ceMDeNM (fig. 7e) and mainly to the free MD (fig. 7d).

**Figure 7.**
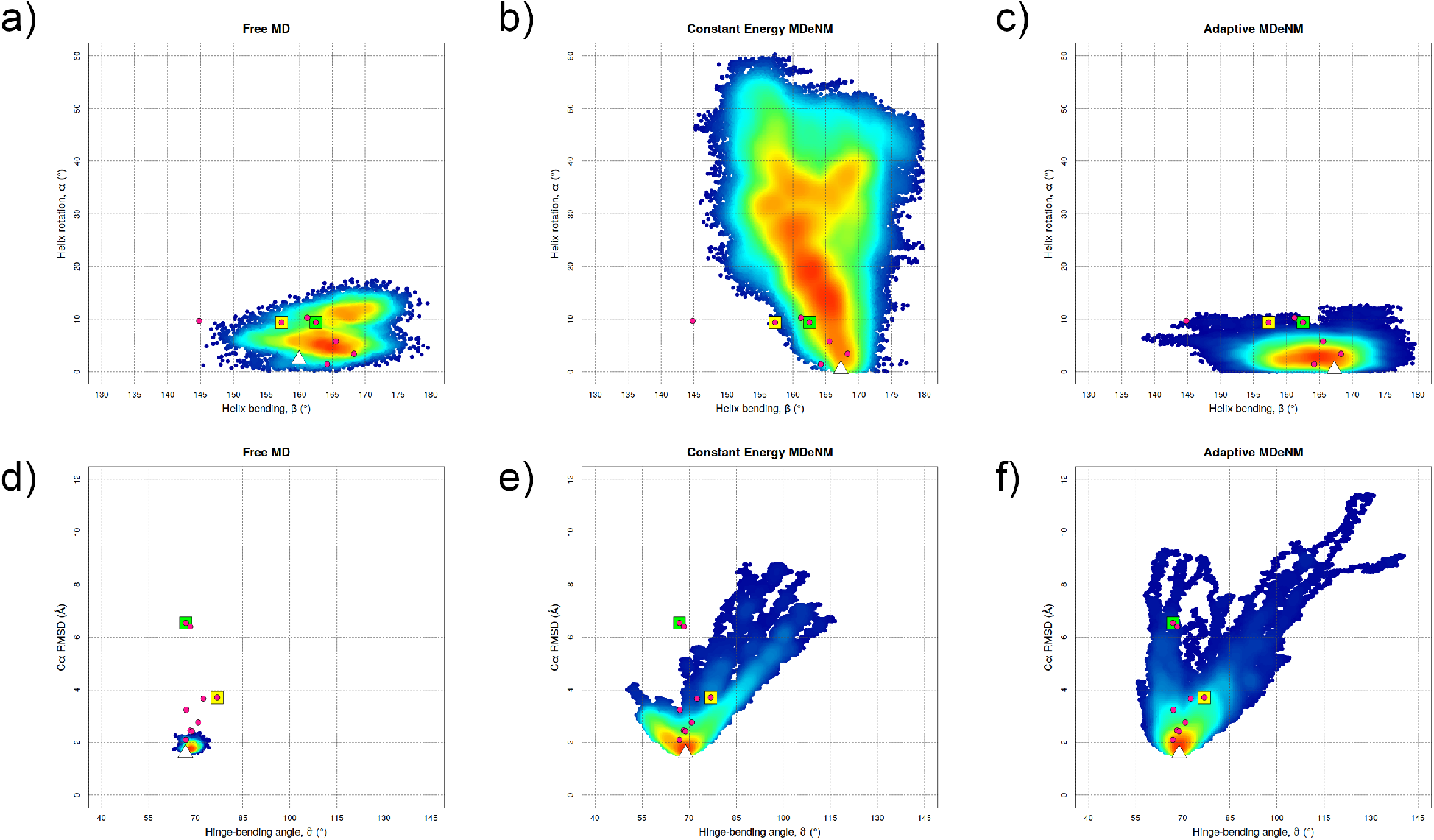
Adaptive MDeNM takes into account structural constraints imposed by the medium. The bending and rotation of the TM helix indicate its orientation inside the membrane. **a)** Free MD present a good sampling of the experimental structure. **b)** However, ceMDeNM showed much larger rotation of the TM helix, indicating a excessive motion from the computed NMs. **c)** On the other hand, aMDeNM presented a similar pattern than the free MD, in accordance with the experimental data. Considering the globular portion of MGT, **d)** the sampling during the free MD was really poor, while **e)** ceMDeNM significantly improved the visited conformations but not as far as **f)** aMDeNM. Taken together, this data shows that aMDeNM take natural constraints imposed by the structure and the environment into account during the conformational sampling.

The correction in in the excitation vectors in MGT is an effect of constraints imposed by the system environment itself. As the modes are computed in a homogeneous dielectric medium, some distortions may be expected in large-scale motions of transmembrane proteins^24^. We observe, indeed, the system is adaptively changing the exploratory direction along MDeNM as a result of the spatial restraint imposed by the membrane. It is not the first time that this effect is observed. The application of an artificial potential that mimes a membrane bilayer during NMA calculation was applied to glutamate transporter and the results presented illustrate that the membrane actually does have an effect in the protein motion^69^. Here, we demonstrate this effect at the atomic level through a method that couples NMA and MD, which allow the comprehensive study of the transition path between the protein’s macrostates.

Here we presented new implementations on MDeNM method that automatically controls the energy injection and take the natural constraints imposed by the structure and the environment into account during protein conformational sampling, which prevent structural distortions all along the simulation. The conformations explored by other enhanced sampling methods, such as REMD, may depict non-physical situations in certain conditions due to changes in energy barriers promoted by the modified temperature^70^. These unrealistic effects can also be observed in collective variable methods, like TMD^71^, in which the lowest energy pathway is not necessarily followed and trajectories may cross large energy barriers undergoing pathways that are inaccessible at room temperature. Preserve the proteins structural features is thus crucial to study its function, since it has been shown to play an essential role on the conformational exploration in equilibrium dynamics^72,73^. Large scale functional motions such as domain shifts are usually characterized by a combination of low-frequency normal modes^62,74^ or PCA^75^ in atomistic level, or even by simplified protein models like ENM^76^ or ANM^77^. Normal modes describe the vibrational spectrum of proteins in a given energetic minimum. Due to the stochasticity of thermal motions, NM eigenvectors move away from the original directions when used to displace the protein, since the structure evolves into other potential energy wells. Therefore, the displacement along the modes is valid for small distances, but the displacement along greater distances may deform the structure of the protein if no care is taken^67^. Many methods were recently developed to took advantage of low-frequency motions to accelerate molecular dynamics simulations and enhance protein sampling. Among then, we discussed ClustENM^26^, ACM^22^, ACM-PCA^66^, NMA-ITS^68^, Saldaño^32^ that also update the excitation direction along the simulation, such as aMDeNM. These methods present similar algorithms but a few different approaches on how to operate the sampling. Nonetheless, rather biasing the potential^68^ or guiding the sampling^26,32^, all of them present the recalculation of new excitation vectors. Our approach differs for there is no need to recompute the normal modes during the simulation or any bias in the potential terms whatsoever. When our algorithm evaluates the displacement along the excitation vector and determines that it is time to update the direction, it is done with respect to the system’s structural and dynamical features. Low and medium frequencies are intrinsically activated during this process, therefore, we do not need to renew the NM input set. Thus, sampling is adaptively updated according to energy barriers, avoiding excessive distortions and conformations physically unattainable at a given temperature.

## 4 CONCLUSIONS

We presented new implementations on MDeNM method that automatically controls the energy injection and take the natural constraints imposed by the structure and the environment into account during protein conformational sampling, which prevent structural distortions all along the simulation.

Normal modes describe the vibrational spectrum of proteins in a given energetic minimum. Due to the stochasticity of thermal motions, NM eigenvectors move away from the original directions when used to displace the protein, since the structure evolves into other potential energy wells. Therefore, the displacement along the modes is valid for small distances, but the displacement along greater distances may deform the structure of the protein if no care is taken. The advantage of this methodology is to adaptively change the direction used to displace the system, taking into account the structural and energetic constraints imposed by the system itself and the medium, which allows the system to explore new pathways.

These implementations could accurately take into account structural constraints imposed to the direction explored in several systems, pointing to their applicability for locating and mapping of energy barriers during conformational sampling. Thus, our data reinforce power of MDeNM in plentiful applications, such as protein−protein flexible docking, structural interpretation of experimental data, conformational exploration of large systems at an atomistic level and recognition of transition paths.

