## Supplementary material for "Adaptive collective motions: a hybrid method to improve conformational sampling with molecular dynamics and normal modes"

[†]Laboratoire de Biologie et Pharmacologie Appliquée, École Normale Supérieure Paris-Saclay, 91190, Gif-sur-Yvette, France.

[¥]Programa de Computação Científica, Fundação Oswaldo Cruz, 21040-360, Rio de Janeiro, Brazil.

[§]Inserm U1268 MCTR, CiTCoM UMR 8038 CNRS – Université Paris Cité , Paris, France.

**AUTHOR INFORMATION**

**Corresponding Author**

David PERAHIA

Laboratoire de Biologie et Pharmacologie Appliquée (LBPA)

École Normale Supérieure Paris Saclay

4 Av. des Sciences

91190 Gif-sur-Yvette, France

Pedro Túlio de RESENDE-LARA

Department of Medical Genetics and Genomic Medicine

School of Medical Sciences , University of Campinas

Rua Tessália Vieira de Camargo, 126 – Campinas, 13083-887, Brazil

**KEYWORDS** Protein dynamics; Conformational sampling; Collective Variables; Normal modes analysis; Molecular dynamics.

### SUPPLEMENTARY MATERIAL

#### 1 MONITORING THE KINETIC ENERGY INJECTION

To demonstrate the importance of controlling the kinetic energy injection, we compared two sets runs, original and ceMDeNM. Each trajectory consisted of 50 ps and the energy injection was fixed to 0.5 kcal along the hinge-bending motion characterized by NM 7 (each set consisting of 3 independent replicas). We observed the fast and cumulative increase in the kinetic energy along the ***Q*** vector in original MDeNM (black curve, supp. fig. 1a). This is due to the fact that the protein does not have enough time to relax and dissipate the additional energy. On the other hand, the excitation energy was maintained at the desired level in ceMDeNM (red curve, supp. fig. 1a). Since mode 7 describes the hinge bending motion, we computed the Cα RMSD from the PDB 3FI5[1], which depicts a closed configuration of T4L (upper inset panel, supp. fig. 1a). We observed that the closing motion is faster in the original approach. However, the RMSD quickly increases after reaching the minimum due to structural distortions (supp. fig. 1b), caused by undissipated energy cumulated during the simulation. Oppositely, ceMDeNM slowly closes the structure without distorting it (supp. fig. 1c). We also computed the Cα RMSD from the initial position (lower inset panel, supp. fig. 1a) and observed that original MDeNM shows a faster increase when compared to ceMDeNM, that increases gradually. Comparing the final structures of MDeNM approaches to the 3FI5 conformation, it is clear that controlling the energy injection presented effective in put less energy into the system and, therefore, obtain better structures at the end of the simulation (supp. fig. 1d).

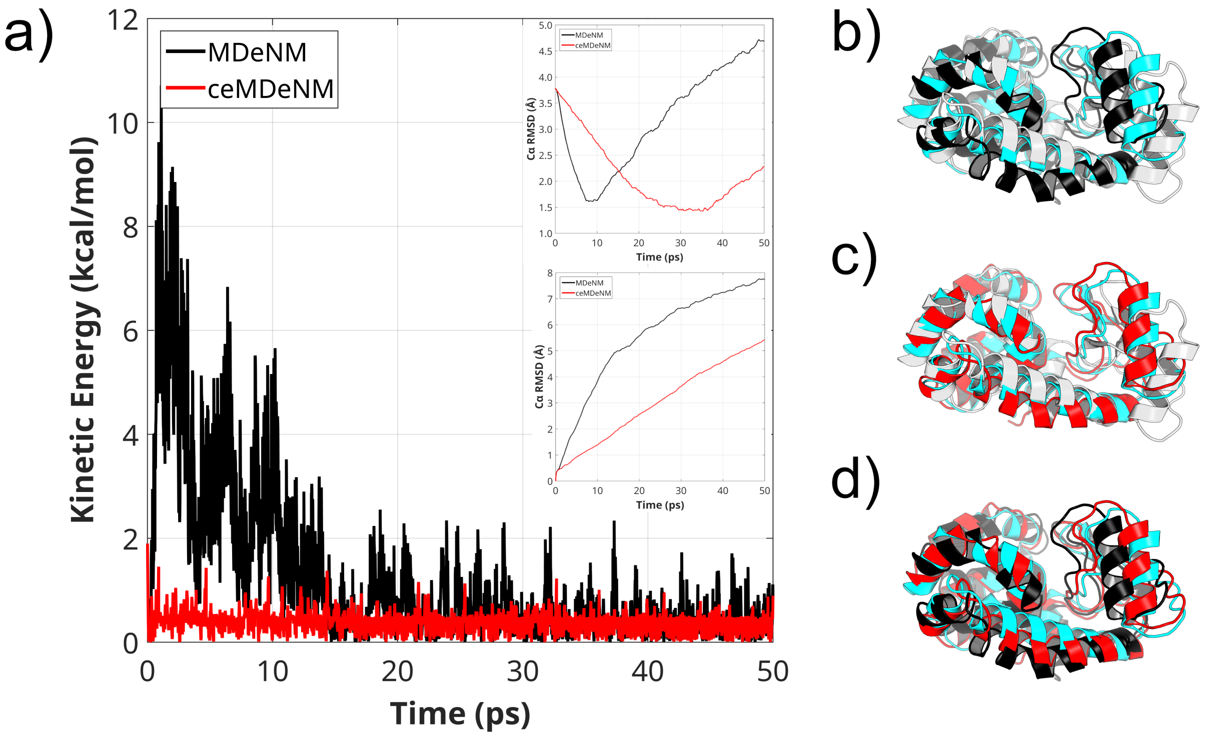


**Supplementary Figure 1. Monitoring the kinetic energy injection along NM 7 in original and constant energy MDeNM. a)** Considering an energy injection of 0.500 kcal/mol every 0.2 ps, the energy injection was evaluated by obtaining the projection of the velocity vector onto the normal mode 7. In ceMDeNM, the energy injection was maintained in a stable plateau around 0.500 kcal/mol during the whole simulations while a great variation was observed in those in which the control was absent. **b)** Initial (light gray) and final (black) conformation of original MDeNM run compared with the crystallographic structure 3FI5 (cyan). We noticed that the structure excessively close without the energy control. **c)** Initial (light gray) and final (red) conformation of ceMDeNM simulation compared with the crystallographic structure 3FI5 (cyan). As previously described, the energy control avoid excessive energy injection, which prevents structural distortions, resulting in a T4L closed conformation with good agreement with experimental data. **d)** Final conformation of original MDeNM (black) and ceMDeNM (red) compared with the crystallographic structure 3FI5 (cyan).


3

### 2 UPDATE OF EXCITATION DIRECTION

As described in the main text, the correction of excitation direction depends on two distinct parameters: i) a given root-mean square displacement value ($\ell$) along the excitation vector $\boldsymbol{Q}$ that describes the effective displacement of the system along this direction; and ii) the deviation of the effective displacement vector from the theoretical one ($cos\ \alpha$). The former dictates to the algorithm the periodicity of evaluation and the latter is the cutoff angle to accept or reject the new excitation direction. In order to evaluate these two parameters, we analyzed the hinge bending motion of the T4 lysozyme, computing the Cα RMSD from the closed configuration represented by PDB 3FI5. The screening of several values of $\ell$ is illustrated in supplementary figure 2 and supplementary table 1. Following the trajectory until the $\ell$ value is reached, no significant difference in RMSD is observed (supp. fig. 2a). However, by projecting these trajectories onto the excitation vector $\boldsymbol{Q}$, we obtain scalar values ranging from 0.29 to 0.63 (supp. table 1).

Supplementary Table 1. Evaluation of the direction deviation from vector Q in different displacements.

| $\ell$ | 0.30 | 0.35 | 0.40 | 0.45 | 0.50 | 0.55 | 0.60 | 0.65 | 0.70 |
|---|---|---|---|---|---|---|---|---|---|
| **$cos\ \alpha$** | 0.29 | 0.40 | 0.46 | 0.48 | 0.47 | 0.52 | 0.57 | 0.63 | 0.63 |

Linear regression shows that $\ell$ and $cos\ \alpha$ are not independent ($p < 0.00001$, data not shown), meaning that an optimized combination of both is necessary to effectively change the excitation direction. Hence, we set $cos\ \alpha$=0.50 in the following essays, which is approximately the average of the sampled values and tested the several $\ell$ values. We then observed the hinge-bending trajectory until the first update takes place, that is, both screened $\ell$ and $cos\ \alpha$ thresholds are reached (supp. fig. 2b). It is interesting to note that small $\ell$ values tend to change the excitation direction earlier than greater ones. In addition, compared to the original MDeNM (black curve), small $\ell$ values entail a direction change fairly before reaching the minimum RMSD value (the


4

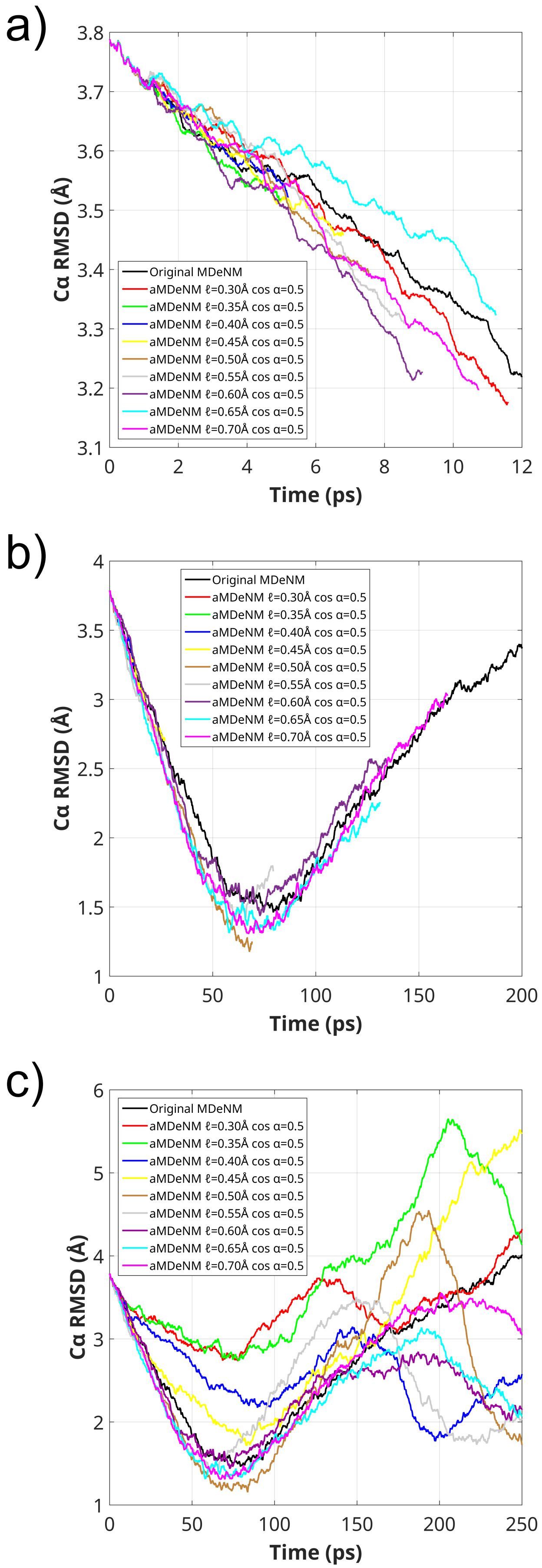

a)
Cα RMSD (Å)
Time (ps)
Original MDeNM
aMDeNM ℓ=0.30Å cos α=0.5
aMDeNM ℓ=0.35Å cos α=0.5
aMDeNM ℓ=0.40Å cos α=0.5
aMDeNM ℓ=0.45Å cos α=0.5
aMDeNM ℓ=0.50Å cos α=0.5
aMDeNM ℓ=0.55Å cos α=0.5
aMDeNM ℓ=0.60Å cos α=0.5
aMDeNM ℓ=0.65Å cos α=0.5
aMDeNM ℓ=0.70Å cos α=0.5
b)
Cα RMSD (Å)
Time (ps)
Original MDeNM
aMDeNM ℓ=0.30Å cos α=0.5
aMDeNM ℓ=0.35Å cos α=0.5
aMDeNM ℓ=0.40Å cos α=0.5
aMDeNM ℓ=0.45Å cos α=0.5
aMDeNM ℓ=0.50Å cos α=0.5
aMDeNM ℓ=0.55Å cos α=0.5
aMDeNM ℓ=0.60Å cos α=0.5
aMDeNM ℓ=0.65Å cos α=0.5
aMDeNM ℓ=0.70Å cos α=0.5
c)
Cα RMSD (Å)
Time (ps)
Original MDeNM
aMDeNM ℓ=0.30Å cos α=0.5
aMDeNM ℓ=0.35Å cos α=0.5
aMDeNM ℓ=0.40Å cos α=0.5
aMDeNM ℓ=0.45Å cos α=0.5
aMDeNM ℓ=0.50Å cos α=0.5
aMDeNM ℓ=0.55Å cos α=0.5
aMDeNM ℓ=0.60Å cos α=0.5
aMDeNM ℓ=0.65Å cos α=0.5
aMDeNM ℓ=0.70Å cos α=0.5



5

**Supplementary Figure 2. Screening of root-mean square displacement ($\ell$) parameter in MDeNM direction correction.** The screening was done by measuring the Cα RMSD from PDB 3FI5. **a)** Stopping the simulations when the $\ell$ threshold is reached, no significant RMSD difference is observed. **b)** Fixing $cos\ \alpha$=0.50 and varying $\ell$, we observed the RMSD curve until the first update in the excitation direction. Comparing to the closed conformation (3FI5), smallest $\ell$ values (red and green curves) updates the direction too soon and for the largest ones (cyan and magenta curves) this is done too late. **c)** RMSD curve for the whole MDeNM simulation (250 ps). This panel illustrates how soon the correction deviates the trajectory from the closing motion for small $\ell$ values (red and green curves). On the other hand, when the threshold was high (cyan and, mainly, magenta curves) the correction was insufficient to change the trajectory, maintaining the excessive closing motion. The threshold of $\ell$=0.50 Å (brown curve) presented the greatest variation in RMSD between the open and closed conformations without structural deformations though. The increase in RMSD after the closing motion was observed once the excitation direction considerably changed, showing that the relaxation pathway was successfully found. The data presented in this figure is the average of 3 independent replicas of each simulation.

closest possible to the 3FI5 configuration) while the greatest $\ell$ values shows almost no difference from the original MDeNM. This indicated that the $\ell$ value should be neither too small (in which the excitation direction is rapidly discarded) nor too large (since the directions will not be correctly updated). We notice that in most of simulations the structure is trapped either in a macrostate or in the transition state (supp. fig. 2c). For small $\ell$ values such as 0.30 Å and 0.35 Å (red and green, respectively), it was not possible to reach the closed state. On the other hand, for 0.55 Å and 0.60 Å (gray and purple, respectively), once the structure reached the closed state, it was trapped. There was almost no direction change for the highest rd values (cyan and magenta). Between $\ell$=0.45 Å or $\ell$=0.50 Å (yellow and brown, respectively), the latter displayed the best results. In these replicas, T4L goes further into the closing motion, exploring very different directions (retaining the secondary structure) while also relaxing and returning to the previous path. Therefore, the chosen $\ell$ and $cos\ \alpha$ values are 0.50 Å and 0.50, respectively.

### 3 SECONDARY STRUCTURE

In addition to the internal measurements presented in the main text, we evaluated the secondary structure conservation along the MDeNM trajectories. The concern about structural distortions arises when a method present an extensive sampling such as our methodology. Taking into account the α-helix content of the experimental dataset of T4L (supp. fig. 3a), we observe

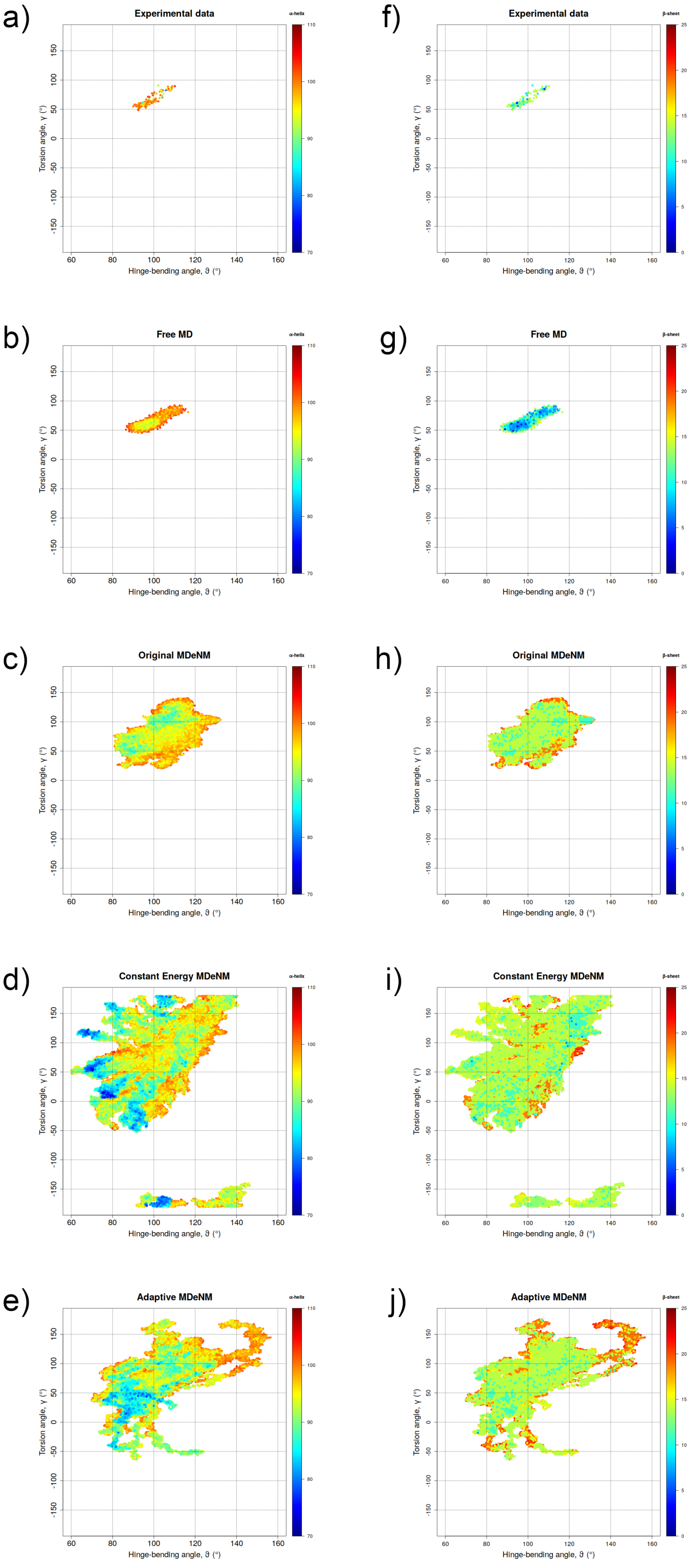

a)
Experimental data
b)
Free MD
c)
Original MDeNM
d)
Constant Energy MDeNM
e)
Adaptive MDeNM
f)
Experimental data
g)
Free MD
h)
Original MDeNM
i)
Constant Energy MDeNM
j)
Adaptive MDeNM
Torsion angle, γ (°)
Hinge-bending angle, ϑ (°)



7

**Supplementary Figure 3. Secondary structure assessment for T4L.** α-Helix content for **a)** experimental data, **b)** free MD, **c)** original MDeNM, **d)** ceMDeNM, and **e)** aMDeNM. While ceMDeNM presented significant loss of helix content, the orignal MDeNM and aMDeNM were more stable. β-Sheet content for **f)** experimental data, **g)** free MD, **h)** original MDeNM, **i)** ceMDeNM, and **j)** aMDeNM. All MDeNM approaches presented similar results for β-sheet content.

that this structural motif was more preserved in the free MD and the original MDeNM (supp. fig. 3b and 3c, respectively) whereas the ceMDeNM presented a relative decrease due to deformations caused by the excessive closing of the protein (supp. fig. 3d). The adaptive approach presented some structures with intermediary values (supp. fig. 3e), but the general features from the experimental structures were also present. Experimental data shows the majority of structures presenting approximately 12 residues in β-sheet (supp. fig. 3f). While the free MD shows some loss of residues in β-sheet (supp. fig. 3g), original MDeNM, ceMDeNM and aMDeNM present similar content proportion of this motif (supp. fig. 3h, 3i and 3j, respectively). All MDeNM approaches also present conformations with mode than 15 residues in β-sheet, indicating the favoring of structuration during the exploration of large scale motions.

CaM structures present a very heterogeneous pattern concerning α-helix content (supp. fig. 4a). This is due mainly to the great flexibility of the interdomain linker, that can be assessed as helix of completely disordered. As expected, since it can not reproduce the several possible states of CaM, α-helix content was poorly represented by the free MD, original MDeNM and ceMDeNM simulations (supp. fig. 4b, 4c and 4d, respectively). We observed an over sampling of well structured conformations, while the ones with less helical content remained unexplored. The updating of the excitation vector allowed the protein to follow different paths and, therefore, change its secondary structure in aMDeNM simulations (supp. fig. 4e). This result is in agreement to the RMSD values presented in the main text, showing that aMDeNM reaches broader sampling considering all experimental structure observed. The β-sheet content results are very similar those presented to α-helix, i. e., the experimental dataset present a heterogeneous


8

pattern (supp. fig. 4f) that are not fully reproduced by free MD, original MDeNM and ceMDeNM (supp. fig. 4g, 4h and 4i, respectively). Again, the adaptive MDeNM presented the closest results from those observed in experimental structures (supp. fig. 4j).


9

114

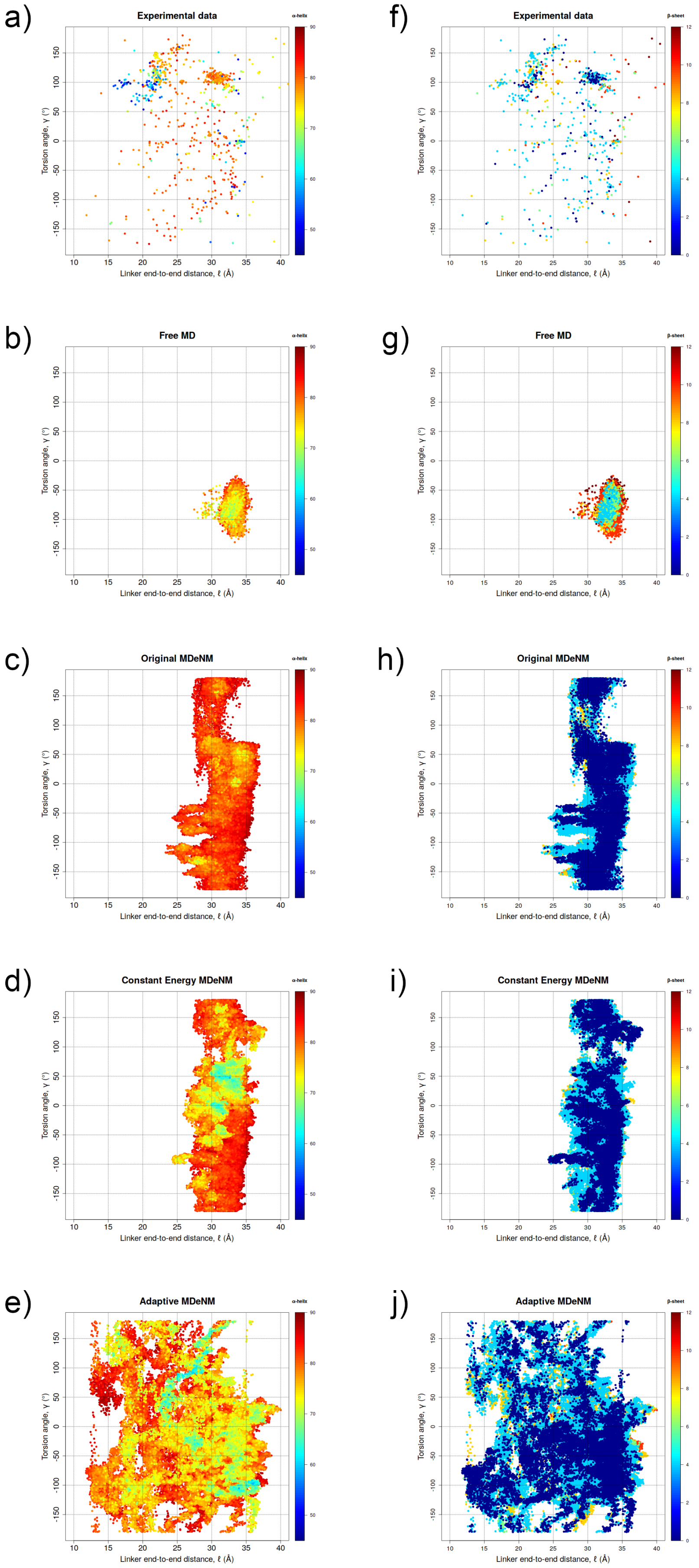

a)
Experimental data
b)
Free MD
c)
Original MDeNM
d)
Constant Energy MDeNM
e)
Adaptive MDeNM
f)
Experimental data
g)
Free MD
h)
Original MDeNM
i)
Constant Energy MDeNM
j)
Adaptive MDeNM
α-helix
β-sheet
Torsion angle, γ (°)
Linker end-to-end distance, ℓ (Å)



10

**Supplementary Figure 4. Secondary structure assessment for CaM.** α-Helix content for **a)** experimental data, **b)** free MD, **c)** original MDeNM, **d)** ceMDeNM, and **e)** aMDeNM. β-Sheet content for **f)** experimental data, **g)** free MD, **h)** original MDeNM, **i)** ceMDeNM, and **j)** aMDeNM. Adaptive MDeNM presented good agreement with the experimental dataset while the other approaches did not properly reproduced the experimental profile.
